# Diverse genome organization following 13 independent mesopolyploid events in Brassicaceae contrasts with convergent patterns of gene retention

**DOI:** 10.1101/120048

**Authors:** Terezie Mandáková, Zheng Li, Michael S. Barker, Martin A. Lysak

## Abstract

Hybridization and polyploidy followed by genome-wide diploidization significantly impacted the diversification of land plants. The ancient At-*α* whole-genome duplication (WGD) preceded the diversification of crucifers (Brassicaceae). Some genera and tribes also experienced younger, mesopolyploid WGDs concealed by subsequent genome diploidization. Here we tested if multiple base chromosome numbers originated due to genome diploidization after independent mesopolyploid WGDs and how diploidization impacted post-polyploid gene retention. Sixteen species representing ten Brassicaceae tribes were analyzed by comparative chromosome painting and/or whole-transcriptome analysis of gene age distributions and phylogenetic analyses of gene duplications. Overall, we found evidence for at least 13 independent mesopolyploidies followed by different degrees of diploidization across the Brassicaceae. New mesotetraploid events were uncovered for tribes Anastaticeae, Iberideae and Schizopetaleae, and mesohexaploid WGDs for Cochlearieae and Physarieae. In contrast, we found convergent patterns of gene retention and loss among these independent WGDs. Our combined analyses of Brassicaceae genomic data indicate that the extant chromosome number variation in many plant groups, and especially polybasic but monophyletic taxa, can result from clade-specific genome duplications followed by diploidization. Our observation of parallel gene retention and loss across multiple independent WGDs provides one of the first multi-species tests that post-polyploid genome evolution is predictable.

**Significance statement:** Our data show that multiple base chromosome numbers in some Brassicaceae clades originated due to genome diploidization following multiple independent whole-genome duplications (WGD). The parallel gene retention/loss across independent WGDs and diploidizations provides one of the first tests that post-polyploid genome evolution is predictable.

## Introduction

Polyploidy or whole-genome duplication (WGD) is ubiquitous across the land plants (Wendel 2015). At least 15% of flowering plant speciation events are the result of polyploidy (Wood *et al*. 2009), and the populations of nearly 13% of diploid species harbor unnamed polyploid cytotypes (Barker *et al*. 2016a). Ancient polyploidy has been identified throughout the evolutionary history of vascular plants (Jiao *et al*. 2011; Li *et al*. 2015) and many angiosperm lineages have a polyploid ancestry (Cui *et al*. 2006; Shi *et al*. 2010; Arrigo and Barker 2012; Jiao *et al*. 2014; Cannon *et al*. 2015; McKain *et al*. 2016; Barker *et al*. 2016b). Many analyses suggest that polyploidy has played a significant role in the diversification of flowering plants (e.g. Soltis *et al*. 2009, 2015; Vanneste *et al*. 2014; Edger *et al*. 2015; Tank *et al*. 2015; Soltis and Soltis 2016). Closely linked with polyploidy is a process of diploidization, reverting the duplicated genome into a diploid(-like) genome and concealing WGD events from detection (Wolfe 2001; Barker *et al*. 2012; Wendel 2015; Dodsworth *et al*. 2016; Mandáková *et al*. 2016; Soltis *et al*. 2016). During diploidization duplicated genes or larger chromosomal regions are lost, some paralogues gain a modified or new function (e.g., Adams and Wendel 2005; Conant *et al*. 2014; Douglas *et al*. 2015; Edger *et al*. 2015), and genome size usually decreases due to recombination-driven removal of repetitive sequences (e.g. Woodhouse *et al*. 2014; Soltis *et al*. 2015) or as a consequence of chromosomal rearrangements including descending dysploidy (e.g. Mandáková *et al*. 2010a, b).

Genome evolution following polyploidy has many predictable aspects. Among descendants of even a single polyploidy, some descendant lineages may share nearly ancestral synteny, whereas other taxa experience relatively more rearrangements and losses (Jaillon *et al*. 2007; Tang *et al*. 2008; Schnable *et al*. 2012; de Miguel *et al*. 2015; Murat *et al*. 2014, 2015). One subgenome of a polyploid genome is often retained with more genes and synteny relative to the other subgenome(s) (Freeling *et al*. 2012; Sankoff and Zheng 2012; Schnable *et al*. 2012; Garsmeur *et al*. 2014; Murat *et al*. 2014; Renny-Byfield *et al*. 2015). In addition to biases resulting from differences in parental genomes, gene function may also influence fractionation. Some functional classes of genes are under-represented, whereas others are over-retained following WGD (e.g., Maere *et al*. 2005; Rensing *et al*. 2007; Barker *et al*. 2008; Bekaert *et al*. 2011; Geiser *et al*. 2016; Li *et al*. 2016). The patterns of convergent retention of functional groups of genes following WGDs are generally consistent with the dosage balance hypothesis (Freeling 2009; Birchler and Veitia 2007, 2010, 2012; Edger and Pires 2009). According to this hypothesis, the drive to maintain dosage balance causes dosage-sensitive genes to be consistently over-retained following WGDs but lost following small scale duplications. Numerous studies have found that the biased signatures of gene retention and loss are consistent with this hypothesis from analyses of the descendants of single WGDs (Maere *et al*. 2005; Rensing *et al*. 2007; Barker *et al*. 2008; Shi *et al*., 2010; Bekaert *et al*. 2011; Geiser *et al*. 2016). However, no study has yet evaluated whether convergent gene retention and loss, consistent with the dosage balance hypothesis, occurs among multiple, related species that have independent WGDs.

Since the pioneering studies revealing the paleotetraploid nature of *Arabidopsis* (Arabidopsis Genome Initiative 2000; Vision *et al*. 2000), later described as the Alpha ( *α*) or At-*α* WGD (Bowers *et al*. 2003; Barker *et al*. 2009), several other WGD events post-dating At-*α* have been discovered. These WGDs can be divided broadly as mesopolyploid and neopolyploid events based upon their age (Mandáková *et al*. 2010a; Kagale *et al*. 2014a; Hohmann *et al*. 2015). Neopolyploids may be identified by their elevated chromosome number, genome size, and largely intact (sub)genomes, contributed by (often extant) parental species. Mesopolyploids, on the other hand, are cytologically more difficult to distinguish from “true diploids”, i.e. diploidized paleopolyploids (e.g. *A. thaliana*), as partial diploidization blurs the primary polyploid character of these genomes (e.g., Lysak *et al*. 2005; Mandáková *et al*. 2010a; Wang *et al*. 2011; Geiser *et al*. 2016). Hybridization between diploidized mesopolyploid populations and species may lead to the origin of neoauto- or neoallopolyploid genomes (e.g. *Brassica napus*).

In mesopolyploid clades, re-diploidization is associated with genome rearrangements and descending dysploidies that sometimes generate a broad spectrum of diploid-like chromosome numbers (e.g. Lysak *et al*. 2005, 2007; Mandáková *et al*. 2010a; Salse 2012). Thus, diploidization following mesopolyploid events frequently explains the extant chromosome number variation, including intra-genus or intra-tribal variation in base chromosome numbers (i.e. origin of dibasic and polybasic taxa). For instance, three base chromosome numbers in *Brassica* (*x* = 8, 9, and 10) and multiple base numbers in the tribe Brassiceae originated from a WGD followed by independent diploidization (Lysak *et al*. 2005, 2007).

Here we aimed to investigate whether multiple base chromosome numbers of the polybasic Brassicaceae genera and tribes resulted from genome diploidization after independent mesopolyploid WGDs post-dating the family-specific At-*α* paleopolyploidization. We approached this question by employing comparative chromosome painting (CCP) and/or whole-transcriptome analysis in **16** species from ten Brassicaceae tribes (Table 1). To corroborate the CCP inferences of mesopolyploidy we analyzed gene age distributions for evidence of large gene duplication events consistent with polyploidy (Barker *et al*. 2008; Barker *et al*. 2010). We also used a combination of among lineage ortholog divergences and a recently developed phylogenomic approach, MAPS (Li *et al*. 2015), to confirm the phylogenetic placement of the inferred WGDs. We show that several diploid-like Brassicaceae genera and tribes have undergone independent mesotetra- or mesohexaploidizations followed by genetic and genomic diploidization concealing the polyploid history of these taxa and generating the karyological variation. Furthermore, we found convergent patterns of gene retention and loss among the independent mesopolyploid WGDs.

**Table 1.**
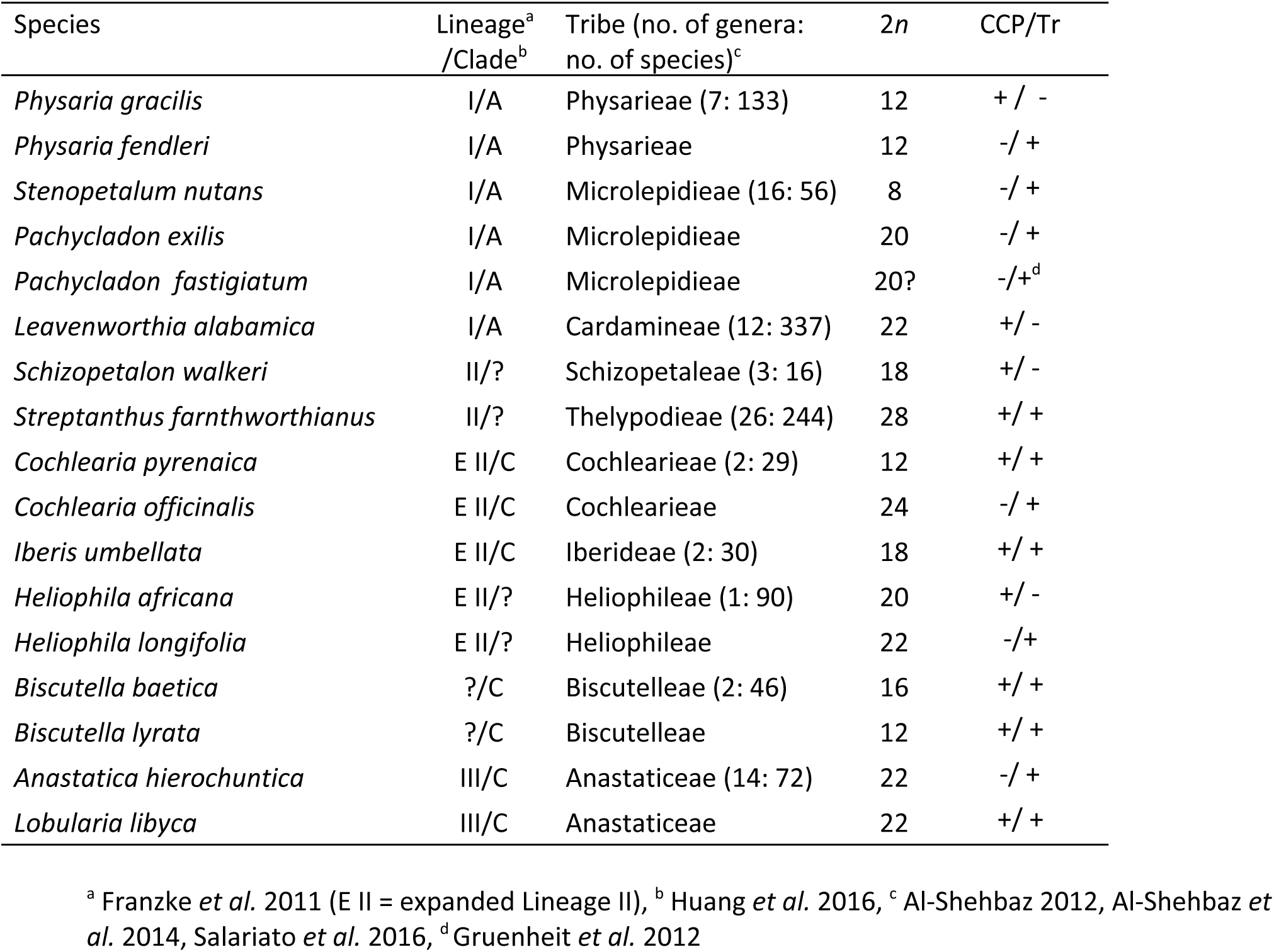
Brassicaceae species analyzed by comparative chromosome painting (CCP) and/or transcriptome sequencing (Tr) in this study.

## Results

### The feasibility of comparative chromosome painting (CCP) in ten Brassicaceae species

We found that CCP with *A. thaliana* BAC contigs is feasible in all analyzed species but with different efficacies. The interpretation of *in situ* hybridization patterns is feasible only if the intensity of hybridized painting probes is sufficiently strong. This is dependent on several factors, such as the phylogenetic distance between *A. thaliana* and each target species, repeat content, or the degree of pachytene chromosome clumping. Moreover, in polyploid genomes, the interpretation of CCP patterns is complicated by diploidization and the associated genomic reorganization. Among 10 analyzed species (Table 1), *L. libyca* had the weakest probe intensity. We used a double concentration of painting probes in this species. The lower fluorescence intensity may reflect the phylogenetic distance between *Arabidopsis* (clade A in Huang *et al*. 2016) and *Lobularia* (clade C). All but three species (*Heliophila africana, Iberis umbellata* and *Physaria gracilis*) demonstrated extensive regions of pericentromeric heterochromatin. This caused clustering of non-homologous centromeres at pachytene. Consequently, interpretation of CCP patterns at proximal chromosomal regions was difficult for most species (Fig. 1).

**Figure 1.**
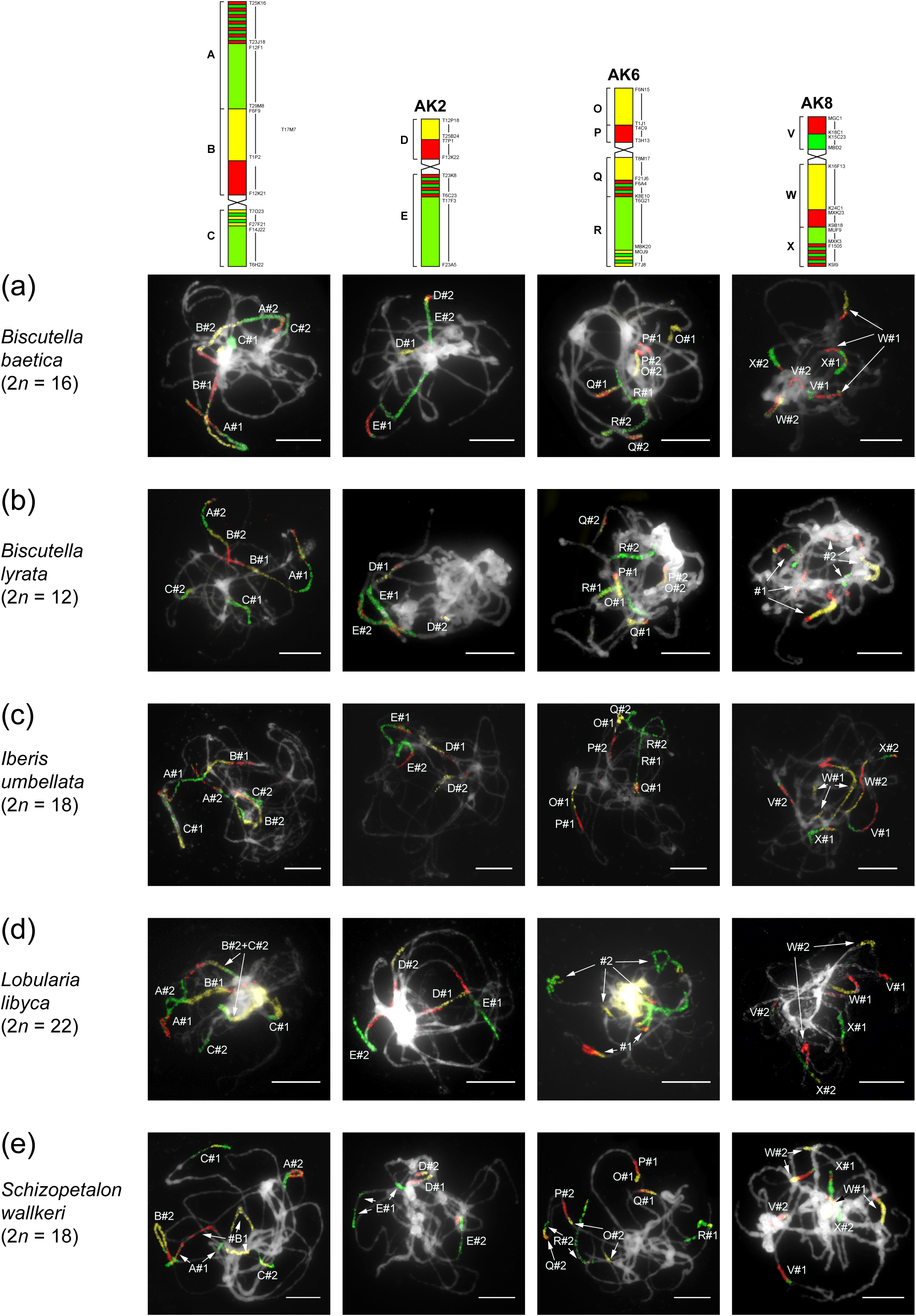
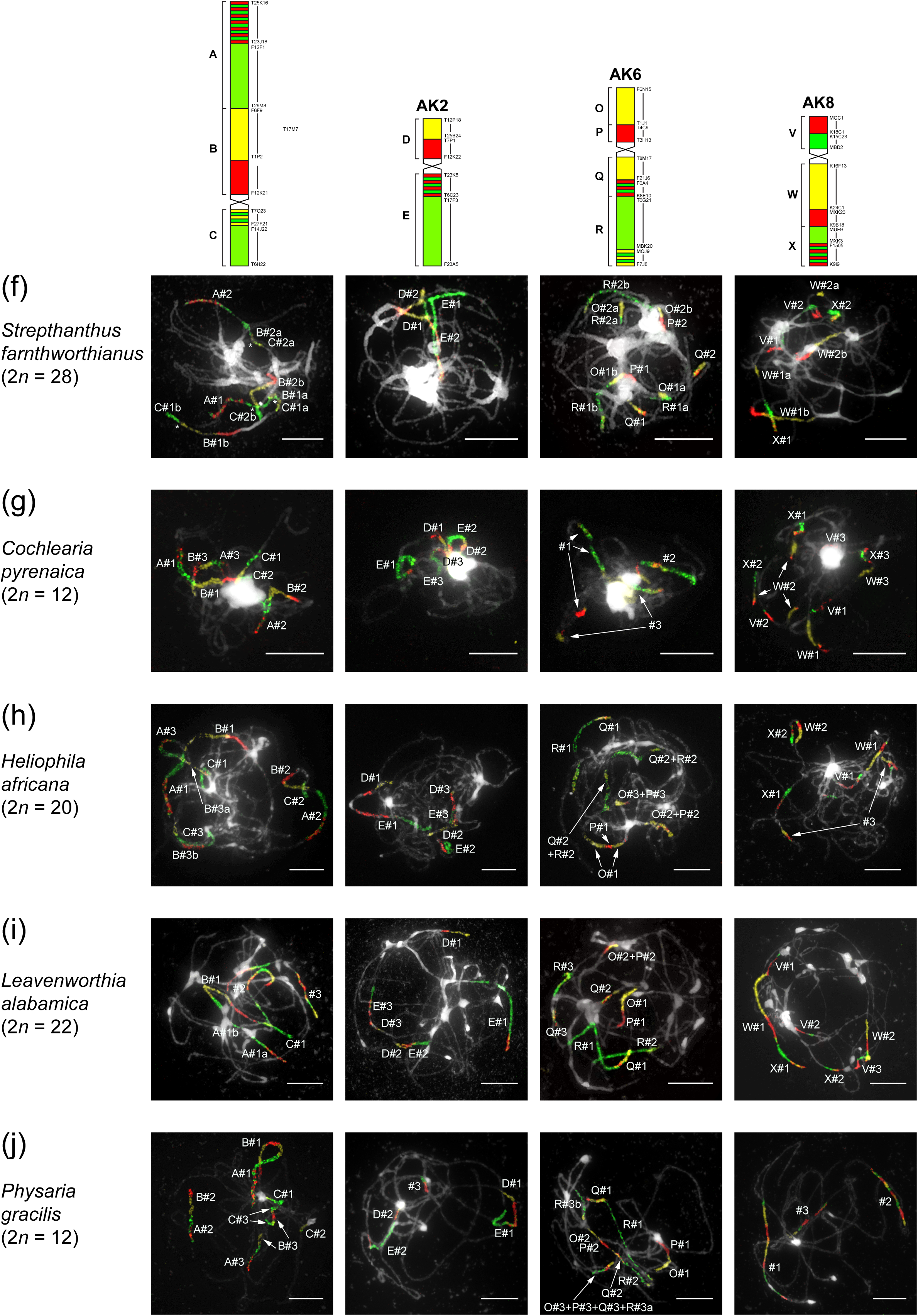
Results of comparative chromosome painting analyses in pachytene (haploid) chromosome complements often Brassicaceae species, (a-f) Mesotetraploid species, (g-j) Mesohexaploid species. The 12 GBs of four ancestral chromosomes were labeled by three different haptens/fluorochromes. Two (a-f) or three (g-j) copies of GBs were distinguished and assigned based on differences in their lenght and fluorescence intensity. All bars, 10 μm.

### New independent mesopolyploid events in Brassicaceae

Previous experimental data (e.g., Lysak *et al*. 2006; Mandáková and Lysak 2008) showed that chromosome-specific BAC contigs of A. *thaliana* hybridize to unique chromosomal regions in Brassicaceae genomes with only the At-*α* WGD. In 10 crucifer species analyzed here, *A. thaliana* BAC contigs representing four ancestral chromosomes (AK1, AK2, AK6 and AK8) with 12 genomic blocks (GB) and ∼48.5 Mb of the *A. thaliana* genome sequence, revealed two or three regions of chromosomal homoeology within the haploid chromosome complements (Fig. 1). The two or three genomic copies of ancestral chromosomes and corresponding GBs were interpreted as resulting from mesotetra- or mesohexaploidization events that occurred in the ancestry of each species after the At-*α* WGD.

### Mesotetraploid genomes

CCP in representatives of Biscutelleae (*Biscutella baetica, 2n* = 16 and *B. lyrata, 2n* = *12*, Fig. 1a and 1b), Iberideae (*Iberis umbellata, 2n* = 18, Fig. 1c), Anastaticeae (*Lobularia libyca, 2n* = 22, Fig. 1d), Schizopetaleae (*Schizopetalon wallkeri, 2n* = 18, Fig. 1e), and Thelypodieae (*Streptanthus farnthworthianus, 2n* = 28, Fig. 1f), revealed two genomic copies of the tested GB associations and unambiguously showed that these species have descended from mesotetraploid ancestors. In two *Biscutella* species and *I. umbellata*, the painted homoeologous GBs differed in length and, in some instances, fluorescence intensity. These differences were more pronounced for longer GBs, whereby a longer and brighter genomic copy was labeled as #1. In *L. libyca, S. wallkeri* and S. *farnthworthianus*, the two genomic copies could not be reliably differentiated. CCP patterns in individual species are presented in Fig. 1 a-f and detailed in Supporting Information, Data S1.

### Mesohexaploid genomes

In representatives of Cardamineae (*Leavenworthia alabamica, 2n* = *22*, Fig. 1i), Cochlearieae (*Cochlearia pyrenaica, 2n* = 12, Fig. 1g), Heliophileae (*Heliophila africana, 2n* = 20, Fig. 1h) and Physarieae (*Physaria gracilis, 2n* = 12, Fig. 1j), CCP uncovered three copies of the GBs that comprise the four ancestral chromosomes. This level of duplication suggests two rounds of WGD in short succession in these species. For simplicity, these are referred to as whole-genome triplication (WGT) events. In all species, triplicated GBs differed significantly in length and, in some instances, fluorescence intensity. This difference was more pronounced in the case of longer GBs (the longest and usually brightest copy always labeled as #1, whereas the shortest and usually the weakest copy always labeled as #3). CCP patterns in individual species are presented in Fig. 1g-j and detailed in Supporting Information, Data S1.

### Transcriptome analyses of mesopolyploidies

Our analyses of transcriptomes from 13 representative Brassicaceae species (Table 1) identified numerous mesopolyploidies and corroborated our cytogenetic results. For the 12 newly sequenced and one previously available (Gruenheit *et al*. 2012) Brassicaceae transcriptomes, we assembled a mean of 40,427 unigenes with an average total assembly size of 29.5 Mb **(Table S2)**. Analyses of gene age distributions found peaks of gene duplication consistent with WGDs in most taxa. Mixture models identified two or three significant normal distributions within the Ks plot of gene duplications for each species (Table 2). These included visually well distinguished peaks in the gene age distributions of *Anastatica, Biscutella, Cochlearia, Heliophila, Pachycladon, Physaria*, and *Stenopetalum*. The gene age distribution in *Lobularia* was relatively noisy, possibly an artifact of its relatively small sample size or potentially indicative of a more complex history of duplication and hybridization (Gaut and Doebley 1997; Barker et al. 2008; Doyle and Egan 2010). Younger inferred WGD peaks in *Iberis* and *Streptanthus* were less distinguished from the initial peak of gene duplications (Fig. 2 and **Fig. S1)**. However, the relatively large number of duplications in the early age distribution is consistent with a young mesopolyploidy in their histories.

**Table 2.**
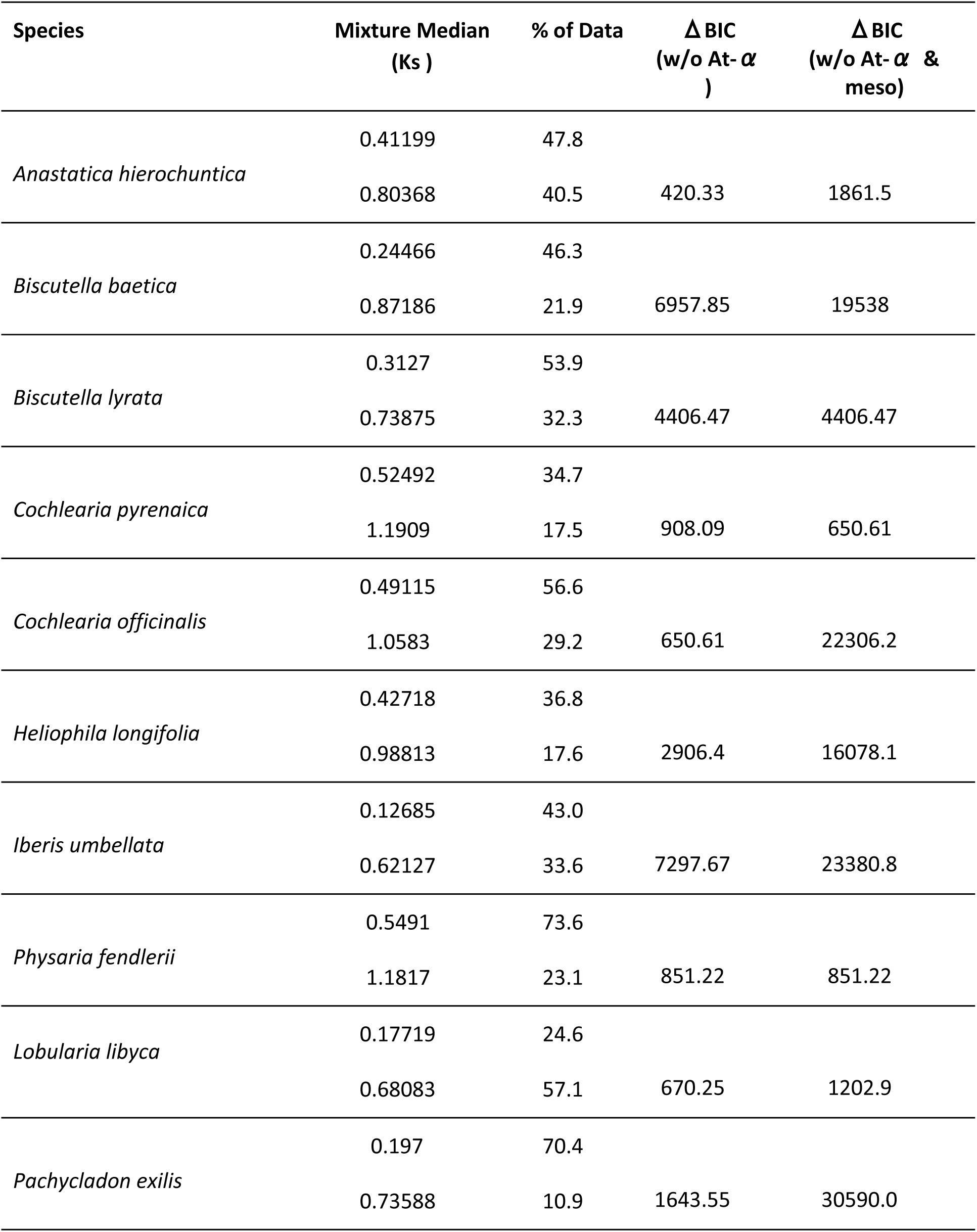
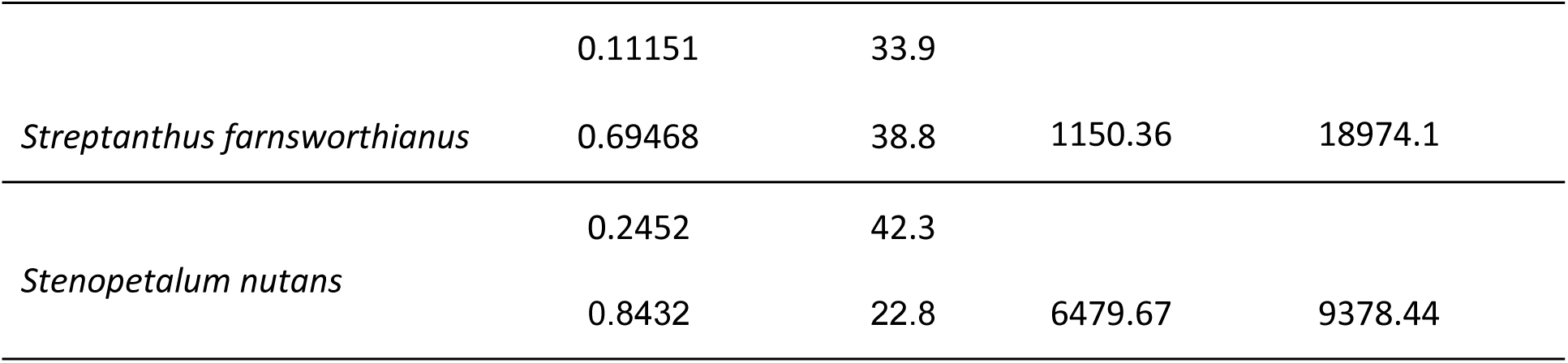
Summary statistics of mixture model distributions inferred to represent the At-*α* WGD and mesopolyploidies across the Brassicaceae taxa analyzed.

**Figure 2.**
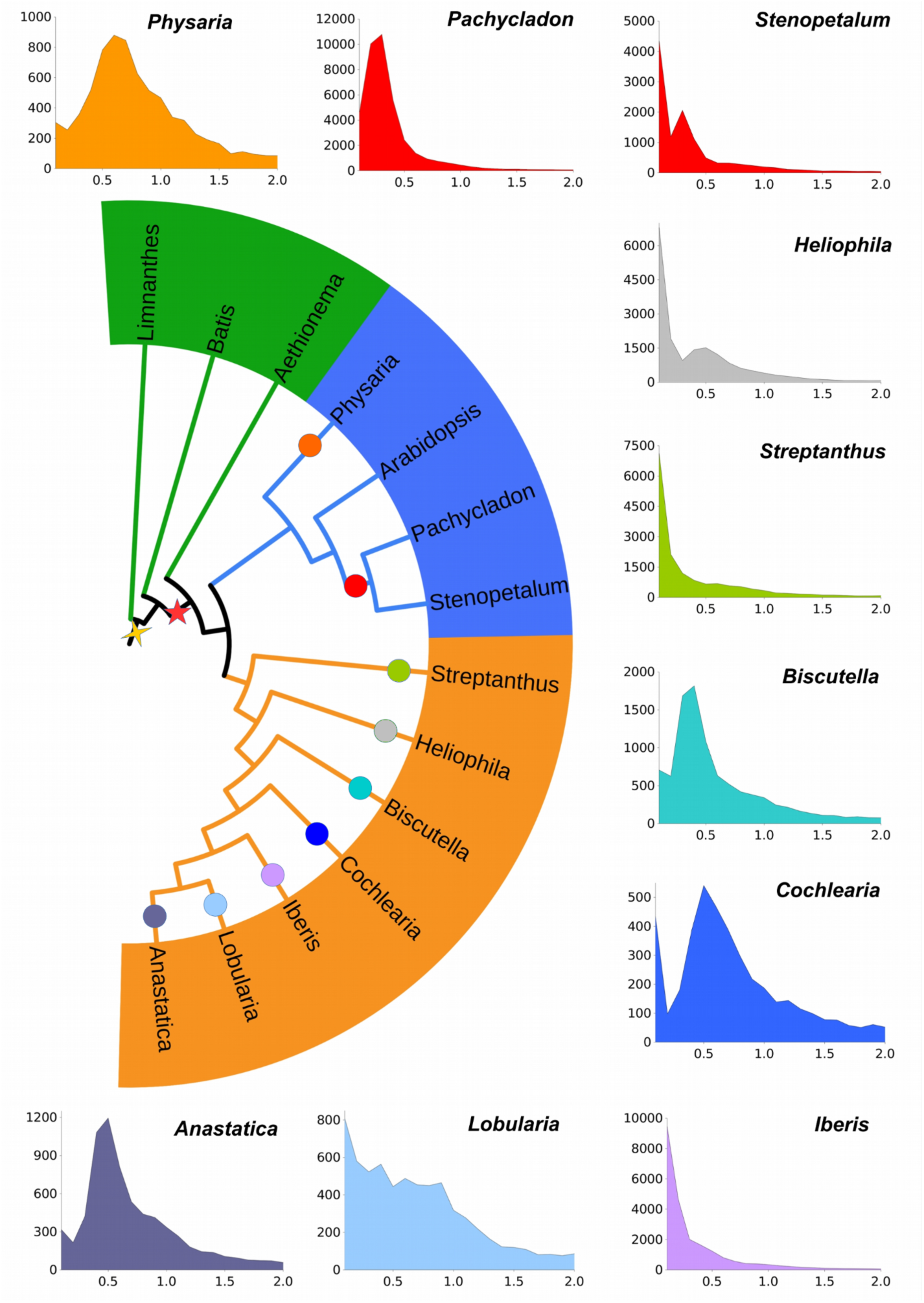
Phylogenetic placement of the nine mesopolyploidies in the Brassicaceae identified by transcriptome analyses. Circles correspond to the inferred location of mesopolyploid events, colors match with 10 Ks plots. Red and yellow stars indicate the inferred Brassicaceae-specific At-*α* and core Brassicales-specific At-*β* paleopolyploidies, respectively. Colors in the simplified phylogeny correspond to broadly defined Lineage I (blue), expanded Lineage II (orange) and Aethionemeae/Brassicales (green).

Each species also contained an older, more fractionated peak located at Ks ∼0.7-1.2. These older peaks likely correspond to the shared At-*α* WGD in the ancestry of all Brassicaceae, whereas the younger and more prominent peaks represent recent mesopolyploidy. The Δ BIC values for models with and without inferred polyploid peaks were large and supported our inference of WGDs across all species (Table 2).

To place these inferred WGDs on the Brassicaceae phylogeny and determine the number of independent WGDs, we used a combination of ortholog divergence (Barker *et al*. 2008, 2009, 2010) and MAPS analyses (Li *et al*. 2015). A comparison of the synonymous divergence of WGD paralogs relative to the divergence between orthologs of species may be used to assess whether a WGD occurred before or after lineage divergence. If the WGD paralog divergence is older than the ortholog divergence, then the WGD likely occurred in a common ancestor of the taxa. Similarly, if the WGD paralog divergence is younger than the ortholog divergence, then the WGD(s) occurred after lineage divergence. In our analyses, the ortholog divergences between genera were generally older than the mesopolyploid Ks divergences and support independent WGDs in *Anastatica, Heliophila, Iberis, Lobularia, Physaria*, and *Streptanthus* (Table 3). However, we found that the ortholog divergence between *Pachycladon* and *Stenopetalum* was more recent than the paralog divergence of their inferred mesopolyploidy, consistent with a WGD in their common ancestor. We also used a recently developed phylogenomic approach, MAPS, to test the phylogenetic placement of these mesopolyploidies. Multi-species gene tree analyses with MAPS supported the ortholog divergence results and found no evidence that the inferred mesopolyploidies in *Anastatica, Heliophila, Iberis, Lobularia, Physaria*, and *Streptanthus* were shared with other taxa. Consistent with independent WGDs within each lineage rather than a shared WGD(s) among genera, the fraction of shared gene duplications in each of these MAPS analyses was low across the analyses for nodes within the Brassicaceae (Fig. 3; **Table S3**). Notably we recovered significant increases in shared gene families for the At-*α* WGD. Overall, our divergence and phylogenomic analyses consistently supported independent mesopolyploidies in the ancestry of *Anastatica, Heliophila, Iberis, Lobularia, Physaria*, and *Streptanthus*.

**Table 3.** Summary of ortholog age distribution analyses. Median ortholog divergence is the median Ks value of ortholog divergences of the selected species pair in each analysis.

| **Taxa1** | **Taxa2** | **Median Ortholog Divergence (Ks)** | **No. of Orthologs** |
| --- | --- | --- | --- |
| *Anastatica hierochuntica* | *Lobularia libyca* | 0.436 | 13364 |
| *Iberis umbellata* | *Lobularia libyca* | 0.489 | 13360 |
| *Iberis umbellata* | *Cochlearia officinalis* | 0.599 | 12696 |
| *Lobularia libyca* | *Cochlearia pyrenaica* | 0.541 | 10434 |
| *Cochlearia pyrenaica* | *Biscutella baetica* | 0.546 | 11354 |
| *Iberis umbellata* | *Biscutella baetica* | 0.494 | 14864 |
| *Biscutella baetica* | *Biscutella lyrata* | 0.169 | 17178 |
| *Cochlearia pyrenaica* | *Cochlearia officinalis* | 0.017 | 9909 |
| *Streptanthus farnsworthianus* | *Biscutella lyrata* | 0.398 | 13718 |
| *Heliophila longifolia* | *Streptanthus farnsworthianus* | 0.435 | 13894 |
| *Heliophila longifolia* | *Lobularia libyca* | 0.472 | 13048 |
| *Pachycladon fastigiatum* | *Stenopetalum nutans* | 0.083 | 6942 |
| *Pachycladon exilis* | *Physaria fendlerii* | 0.450 | 14602 |

**Figure 3.**
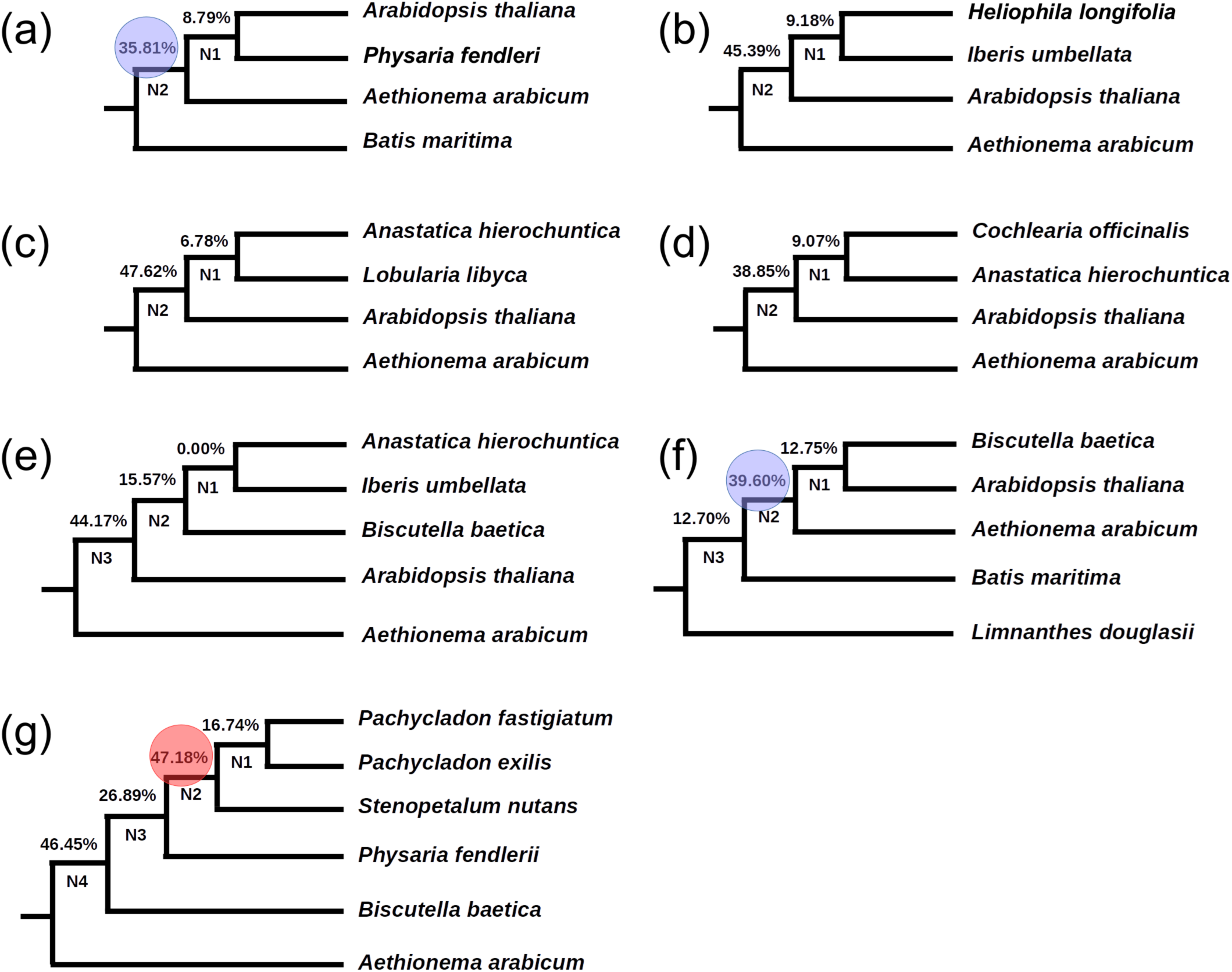
Species trees with gene duplication at each node as inferred by MAPS. The percentage of gene subtrees consistent with the species tree that support a shared duplication is indicated at each node, (a and f) Blue circles highlight inferred Brassicaceae-specific At-*α* paleopolyploid event at the associated nodes, (a-f) The low percentage of gene subtrees at other nodes support independent mesopolyploid events in *Anastatica, Biscutella, Cochlearia, Heliophila, Iberis, Lobularia, Physaria*, and *Streptanthus*. (g) A red circle highlights the inferred mesopolyploidy shared by *Pachycladon* and *Stenopetalum*.

We also found evidence that some of the inferred WGDs occurred in the common ancestor of pairs of genera. Our analyses of *Pachycladon* and *Stenopetalum* indicated that they share a mesopolyploidy. Although the median Ks values of their WGD peaks are significantly different (Ks = 0.197 vs 0.245), the ortholog divergence between *Pachycladon* and *Stenopetalum* was lower and more recent than their mesopolyploid paralog divergence (Table 3). Analyses of substitution rates across 3403 nuclear orthologs indicated that *Stenopetalum* is evolving ∼21% faster than *Pachycladon* relative to *Arabidopsis*. Accounting for this substitution rate heterogeneity aligned the median Ks divergence estimates for the putative shared WGD. The median paralog divergence for the *Stenopetalum* WGD was Ks = 0.245, whereas the median rate corrected paralog divergence for the *Pachycladon* WGD was Ks = 0.238. Consistent with our ortholog divergence and rate analyses, greater than 47% of the gene subtrees in the MAPS analyses supported a shared WGD in the ancestry of *Pachycladon* and *Stenopetalum* (Fig. 3). Similarly, ortholog divergences were much younger than the paralog divergence of inferred mesopolyploidies in the two within genera comparisons in *Biscutella* and *Cochlearia*. Overall, we found evidence for nine independent mesopolyploidies in the ancestry of the analyzed taxa.

Each species also contained a mixture with a median Ks ∼0.7-1.2 that was consistently significant by large Δ BIC values. Previous analyses of the Brassicaceae found that paralogs from the At-*α* WGD diverge in this range of synonymous substitution (Barker *et al*. 2009; Edger *et al*. 2015). Analyses of ortholog divergence indicated that all Brassicaceae lineages diverged after these Ks ∼ 0.7-1.2 WGD peaks (Table 3), consistent with these peaks representing the At-*α* paleopolyploid event. Finally, our MAPS analyses, as expected, indicated that all Brassicaceae share the At-*α* WGD (Fig. 3 a and f). More than 35% of gene subtrees consistent with the species tree supported a shared gene duplication event in the ancestry of all Brassicaceae, consistent with At-*α* The other MAPS analyses did not include a non-Brassicaceae outgroup (Fig. 3 b, c, d, e, and g) and did not directly test the node where At-*α* is established to occur (Barker *et al*. 2009; Edger *et al*. 2015).

### Biased gene retention and loss

Given the numerous mesopolyploidies and karyotypic variation revealed by our analyses, we evaluated the consistency of post-polyploid genome evolution across the diverse Brassicaceae species sampled. To explore the patterns of gene retention and loss following each mesopolyploidy and the At-*α* WGD, we annotated each transcriptome using the *Arabidopsis* gene ontologies. For each species, we used the mixture model Ks ranges for each WGD to identify paralogs from the mesopolyploidies and the At-*α* WGD. The GO categories for these mesopolyploidy and At-*α* WGD paralogs were then analyzed using a simulated chi-squared test to identify significantly over- and under-retained GO categories among these WGD paralogs. We also used a hierarchical clustering approach to group WGD GO category profiles by similarity. Consistent with our expectations, we found evidence that the patterns of paralog retention and loss from all nine mesopolyploidies and the At-*α* WGD were significantly biased with respect to GO categories and largely convergent across the independent mesopolyploidies (Fig. 4). Overall, the GO categories retained and lost following all of the WGDs were generally similar in profile, but there were some consistent differences of GO category enrichment based on WGD age. Hierarchical clustering resolved two major clusters of gene retention and loss among the mesopolyploid and At-*α* paralogs. One cluster was largely composed of genes retained or lost from independent mesopolyploidies, whereas the other cluster consisted mostly of genes retained and lost following the shared At-*α* WGD (Fig. 4a). Chi-squared analyses of GO category retention and loss identified many significantly over- and under-retained functional categories of paralogs (Fig. 4b). Paralogs retained from the At-*α* event were significantly enriched across our sampled species for genes associated with other cellular processes, other metabolic processes, other binding, transferase activity, protein metabolism, kinase activity, transcription factor activity, and signal transduction. Many of these GO categories were significantly under-retained among the paralogs in mesopolyploidies. In contrast, the independent mesopolyploidies consistently demonstrated significant over-retention of genes associated with other intracellular components, other cytoplasmic components, other membranes, chloroplast, plastid, extracellular, and electron transport. Similarly, many of these GO categories were significantly under-retained among At-*α* paralogs across our sampled species (Fig. 4).

**Figure 4.**
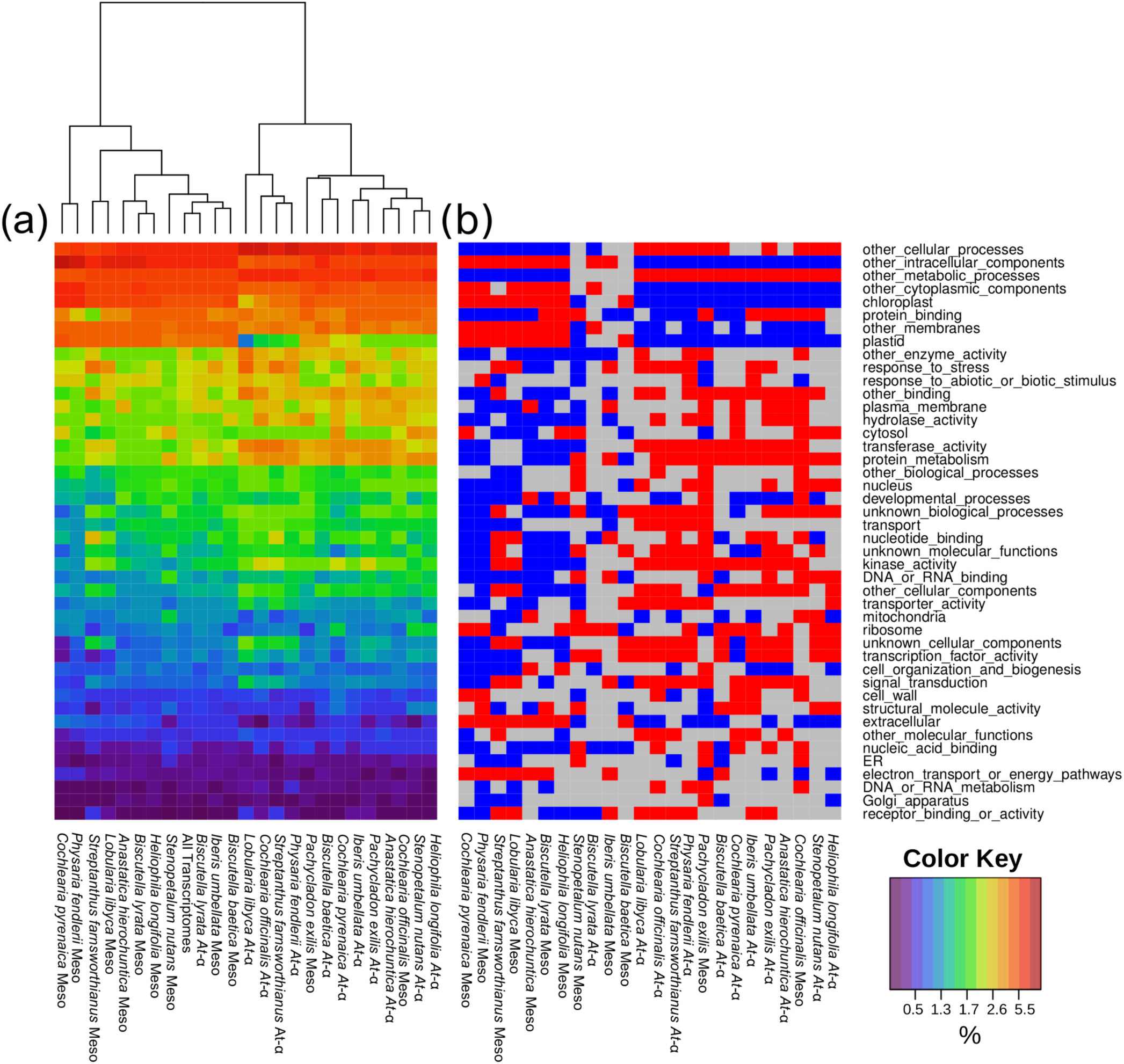
GO annotations of Brassicaceae whole transcriptomes and paleologs. Each column represents the annotated GO categories retained in duplicate following independent mesopolyploidies or the At-*α* WGD from each analyzed species, whereas the rows represent a particular GO category. In (a), colors of the heat map represent the percent of the transcriptome represented by a particular GO category, with red being highest and purple lowest. The overall ranking of GO category rows was determined by the ranking of GO annotations among the total pooled transcriptomes. Hierarchical clustering was used to organize the heatmap columns. In (b), GO categories are identified that are significantly over- (red) or under-retained (blue) among paleologs relative to the total pooled transcriptomes, as determined by residuals from chi-square tests. GO categories with gray boxes in the (b) panel are not represented in significantly different numbers than expected by chance.

Although these contrasting patterns of gene retention and loss between mesopolyploidies and the At-*α* WGD were largely consistent across taxa, there were a few exceptions to this pattern (Fig. 4). The paralogs from the At-*α* event of *Biscutella lyrata* clustered with the mesopolyploid paralogs of most species. However, relatively few GO categories of *B. lyrata’s* At-*α* paralogs were significantly over or under retained, and its mesopolyploid paralogs had a similar pattern of GO categories to most other mesopolyploidies. It appears that *B. lyrata’s* At-*α* paralogs clustered with the mesopolyploidies because it is an outlier rather than sharing great similarity. Low quality tissue or other sample preparation variation for *B. lyrata* may have caused poor sampling of the genes. This is supported by the fact that the differential gene retention and loss in *B. lyrata’s* congener, *B. baetica*, follows the general pattern. In contrast, the retained paralogs from the mesopolyploidies of *Cochlearia officinalis* and *Pachycladon exilis* had a pattern of GO categories similar to the At-*α* paralogs. These mesopolyploid paralogs are not clearly outliers and seem to have a similar pattern of retention to At-*α* paralogs from their own and other species genomes. The pattern of gene retention following the mesopolyploidy in *Pachycladon* was significantly different than gene retention and loss from the same WGD in *Stenopetalum*. Although *Pachycladon’s* nuclear substitution rate is ∼21% slower than *Stenopetalum*, the pattern of gene retention did not appear to reflect a simply slower rate of loss to the mesopolyploid pattern. It may be that a significantly different substitution rate impacts the pattern of genome fractionation, especially if genes are lost by accumulating substitutions. As *Pachycladon* is relatively slowly evolving and C. *officinalis* is a young neomesopolyploid species (2*n* = 4x = 24), duplicated At-*α* paralogs may have not yet undergone gene fractionation comparable with the remaining mesopolyploid species. For C. *officinalis*, this is also supported by gene retention and loss patterns in the mesopolyploid C. *pyrenaica* which follows the general pattern.

## Discussion

### Cytogenetic and transcriptomic evidence of clade-specific mesopolyploid events across Brassicaceae

Here for the first time we combined large-scale comparative chromosome painting and transcriptome analyses to identify and characterize WGD events across the Brassicaceae. Using painting probes for half of the ancestral *n* = 8 chromosome complement we uncovered new mesotetraploidy events putatively specific for Anastaticeae (*Anastatica-* and *Lobularia*-specific WGD), Iberideae and Schizopetaleae, and two new mesohexaploidy events in the ancestry of *Cochlearia* (Cochlearieae) and *Physaria* (Physarieae). Furthermore, we confirmed a mesotetraploidy in *Biscutella* (Biscutelleae, Geiser *et al*. 2016) and Thelypodieae (Burell *et al*. 2011; Kagale et al. 2014a), another event in Microlepidieae (WGD shared by *Pachycladon* and *Stenopetalum;* Mandáková *et al*. 2010a, b), as well as the mesohexaploidy events in *Leavenworthia* (Cardamineae, Haudry *et al*. 2013) and Heliophileae (Mandáková *et al*. 2012).

Although cytogenetic analyses can reveal the existence and nature of mesopolyploidies, it may be difficult to discern if they represent shared or independent ancestral WGDs. This is particularly true if post-polyploidization genome reshuffling obscures either ancestrally shared or genome-specific cytogenetic signatures. Analyses of newly sequenced transcriptomes from ten Brassicaceae genera corroborated our cytogenetic analyses. Our combined single species and phylogenomic approaches found that nearly all of the analyzed genera experienced an independent mesopolyploidy. Overall, our cytogenetic and transcriptomic analyses revealed six new independent mesopolyploidies in the ancestry of polybasic Brassicaceae genera and tribes (Fig. 2 and **Fig. S1**).

### Mesotetraploidy events

CCP and transcriptome analysis unveiled a previously unknown WGD in *Iberis umbellata* (*n* = 9) belonging to 27 *Iberis* species from the rather small tribe Iberideae (2 genera, 30 spp.; Al-Shehbaz 2012). The WGD is most probably shared by the whole genus and various base numbers (*n* = 7, 9, and 11; Warwick and Al-Shehbaz 2006) reflect the different extent of post-mesopolyploid diploidization in *Iberis*.

Another newly described mesotetraploidy occurred in the ancestry of the *S. walkeri* genome (*n* = 9). *Schizopetalon* (10 spp.) together with *Atacama* (1 sp.) and *Mathewsia* (6 spp.) constitute the small tribe Schizopetaleae confined exclusively to central Chile and parts of Peru and Argentina (Salariato *et al*. 2016). More data are needed to determine if the WGD is shared by all of Schizopetaleae and the CES clade (Salariato *et al*. 2016).

By cytogenetic and transcriptomic analysis of *Streptanthus farnthworthianus* (*n* = 14) we demonstrated that this species and most likely the whole Thelypodieae (26 genera, 244 spp.), dominated by species with *n* = 14 and 13, have a neo/meso tetraploid origin based on *n* = 7. CCP analysis suggested that the S. *farnthworthianus* genome has originated by duplication of the Proto-Calepineae Karyotype (PCK, *n* = 7; Mandáková and Lysak 2008). This is in accordance with the inclusion of Thelypodieae into Lineage II (clade B sensu Huang *et al*. 2016). The Thelypodieae-specific WGD was already suspected by Burell *et al*. (2011) from genetic mapping in *Caulanthus amplexicaulis* (*n* = 14) and further corroborated by transcriptome sequencing in *Pringlea antiscorbutica* (*n* = 12) and *Stanleya pinnata* (*n* = 12 or 14) (Kagale *et al*. 2014a). Chromosome number variation across Thelypodieae (*n* = 7, 9, 10, 11, 12, 13, and 14) and our data suggest that the primary neo/mesotetraploid genome had 14 chromosomes and that the lower chromosome counts represent descending dysploidies.

Despite the prevalent low, diploid-like, chromosome numbers (*n* = 4-7, 10) of the endemic Australian/New Zealand crucifers classified as members of Microlepidieae (16 genera, c. 56 species; Heenan *et al*. 2012), the group has descended from allotetraploid ancestors with approximately 16 chromosomes (Mandáková *et al*. 2010a, b). Besides the 15 endemic Australian genera, Microlepidieae contain eleven species of the genus *Pachycladon* endemic to New Zealand (10 spp.) and Tasmania (1 sp.). Unlike the Australian species, *Pachycladon* species show a remarkable stasis of chromosome numbers (all *n* = 10) and ancestral genome structures (Mandáková *et al*. 2010b). The Australian genera and *Pachycladon* species differ in the extent of genome diploidization, whereby the Australian taxa have undergone more extensive post-polyploid genome repatterning relative to ten *Pachycladon* chromosomes built by largely preserved 16 (2 × 8) ancestral chromosomes. Surprisingly, we found that *Pachycladon* and *S. nutans*, a representative of the Australian Microlepidieae taxa, shared the mesotetraploid WGD. Given that they began with the same parental genomes, the extent of diploidization differentiating the two Microlepidieae lineages is likely explained by a much slower tempo of diploidization in *Pachycladon*. Consistent with this hypothesis, our analyses of nuclear substitution rates indicates that the Australian taxa are evolving more than 20% faster than the New Zealand *Pachycladon*. These results indicate that rates of diploidization are correlated with differences in substitution rates. Previous research has found that paleopolyploid genomes of relatively slow evolving taxa, such as *Vitis vinifera* (Cenci *et al*. 2013; Murat *et al*. 2015), are less rearranged than the genomes of faster evolving taxa. These results are the first to demonstrate this pattern in taxa that descend from the same mesopolyploidy that is recent enough to reconstruct their karyotypes.

## Mesohexaploidy events are not rare in the Brassicaceae

Until recently, the only mesohexaploidy known in the Brassicaceae was the WGT detected in *Brassica* and probably shared by the whole Brassiceae (Lysak *et al*. 2005, 2007; Parkin *et al*. 2005). Later, a tribe-specific WGT was reported to occur in the ancestry of the Heliophileae (Mandáková *et al*. 2012) and in the North American *Leavenworthia* (Haudry *et al*. 2013). Here, by uncovering WGT events in Cochlearieae and Physarieae, we show that mesohexaploidy events might be more frequent than initially thought.

*Leavenworthia* (8 spp.) is confined to New World and the available chromosome counts mapped on a phylogenetic tree (Urban and Bailey 2013) suggest that *n* = 11 is a descending dysploidy from *n* = 15, and most likely both chromosome numbers were derived from the primary hexaploid genome with *n* = 24 chromosomes (2*n* = *6x* = 48). Considering that *Leavenworthia* and *Selenia* (5 spp.) form a well defined, monophyletic group, with some species of *Leavenworthia* and *Selenia* co-occurring, it can be assumed that the WGT predates the divergence of two sister genera and that it contributed to diversification and species radiation within this North American Cardamineae subclade.

Physarieae is one of the most species-rich Brassicaceae clades with seven genera harboring c. 133 species. They are primarily distributed in North America, but several species are disjunctly distributed in Argentina and Bolivia (Al-Shehbaz 2012). Physarieae was repeatedly confirmed to be a monophyletic group with two large subclades: DDNLS (5 genera) and PP (*Paysonia* and *Physaria*) (Fuentes-Soriano and Al-Shehbaz 2013). *Paysonia* has only eight species with diploid-like chromosome numbers of *n* = 7, 8 and 9, whereas *Physaria* is a large genus of 106 species with more variable chromosome numbers (*n* = 4, 5, 6, 7, 8, 9, 10, 12, and higher counts; Warwick and Al-Shehbaz 2006). Both transcriptome and cytogenetic data suggest that at least *Physaria* has undergone an ancestral WGT event followed by species radiations and genome diploidization towards the extant spectrum of diploid-like chromosome numbers.

Cochlearieae contains 29 species in only two genera - *Cochlearia* (20 spp.) and *Ionopsidium* (9 spp.) (Al-Shehbaz 2012). We analyzed a species with the lowest chromosome number known in the tribe (*n* = 6) and obtained convincing data indicating that the genus *Cochlearia* has descended from a mesohexaploid ancestor. Subsequent diploidizing reduction of chromosome numbers resulted in the origin of diploid-like chromosome complements, and some of the mesohexaploids formed new allopolyploid species (Koch *et al*. 1998). Interestingly, Kagale *et al*. (2014a) missed the *Cochlearia*-specific WGT by analyzing the neomesohexaploid genome of *C. officinalis* (*n* = 12).

Mandáková *et al*. (2012) by analyzing seven different *Heliophila* species concluded that probably the whole Heliophileae containing more than 90 species, endemic to South Africa and Namibia, experienced a shared WGT. Here, by analyzing two more species by CCP (*H. africana, n* = 10) and transcriptome sequencing (*H. longifolia, n* = 11), we further corroborated our previous conclusion on incidence of this tribe-specific WGT event.

### How outstanding is the number of mesopolyploidies observed in the Brassicaceae?

In total there are 13 mesopolyploid WGDs, including the WGT and WGD specific for Brassiceae and *Orychophragmus* (Lysak *et al*. 2005, 2007), which occurred in the Brassicaceae crown group after the At-*α* paleotetraploidization and after the split of Aethionemeae and the crown group **(Fig. S1)**. Recognizing 49 Brassicaceae tribes (Al-Shehbaz 2012), at least 11 tribes (22%) have a mesopolyploid ancestry. All but one (as there is not a sufficient number of chromosome counts for Schizopetaleae) mesopolyploid tribes are polybasic, and thus, multiple base chromosome numbers in a genus or tribe may indicate an ancient WGD followed by independent diploidizing descending dysploidies. Given our observations in the Brassicaceae, it may be that most polybasic genera of plants have descended from mesopolyploid ancestors. However, most analyses of genomic data from families of plants have not revealed evidence for WGDs in the history of each analyzed genus. For example, genomic analyses that included data from many genera of the Asteraceae (Barker *et al*. 2008; Barker *et al*. 2016b) support numerous ancestral polyploidies, but there is no evidence that every genus experienced an independent ancestral WGD. This is particularly surprising in the light of our present findings and the enormous taxonomic (c. 1,623 genera with 24,700 species, Funk *et al*. 2009) and karyological (Jones 1985) diversity of the Asteraceae. Similar results have been observed in studies of the Poaceae (Jiao *et al*. 2014; McKain *et al*. 2016), Fabaceae (Cannon *et al*. 2015) and Orchidaceae, one of the largest vascular plant families with c. 880 genera and up to 27,000 species where only one paleopolyploid WGD was revealed so far (Cai *et al*. 2015). In light of these comparisons, it seems that the frequency of mesopolyploidy in the Brassicaceae may be a relative outlier. It should be however noted that this conclusion is premature as many analyses of plant genomes have not included such karyotypically variable lineages and more mesopolyploidies may be found as these lineages are sequenced.

Recent transcriptomic analyses found neo/mesopolyploid events in eight species from five Brassicaceae tribes (Kagale *et al*. 2014a). However, five of these WGDs are likely best characterized as neopolyploidies based on their elevated euploid chromosome numbers. These neopolyploids include *Capsella bursa-pastoris* (2*n* = 4*x* = 32; Douglas *et al*. 2015), *Armoracia rusticana* (2*n* = 4*x* = 32), *Draba lactea* (2*n* = 4*x*, 6*x* = 32, 48; Grundt *et al*. 2005), *Lepidium densiflorum* (2*n* = 4*x*= 32; http://brassibase.cos.uni-heidelberg.de/) and *L. meyenii* (2*n* = 8*x* = 64; Quiros *et al*. 1996, Zhang *et al*. 2016). The puzzling chromosome number of *L. sativum* (2*n* = 24) most likely represents diploidization of a neo/mesotetraploid genome (2*n* = 4*x* = 32), as suggested by Graeber *et al*. (2014) and observed in *Cardamine cordifolia* (Mandáková *et al*. 2016).

### Ancestral mesopolyploid genomes and structural diploidization

Polyploid genomes of the Brassicaceae originate through duplication of either Ancestral Crucifer Karyotype (ACK) with *n* = 8 or Proto-Calepineae Karyotype (PCK) with *n* = 7 (Lysak *et al*. 2016). Mesotetraploidy thus yields genomes with 16 or 14 chromosomes, whereas mesohexaploidy results in genomes with 24 or 21 chromosomes (for the sake of simplicity hybridization between *n* = 8 and *n* = 7 genomes is not considered). The degree of diploidization, i.e. the reduction of ancestral chromosome numbers (*n* = 14, 16, 21 or 24) to extant, diploid-like chromosome numbers, appears to differ among the groups of mesopolyploid origin (Fig. 5). The lowest chromosome numbers, and thus the highest degree of structural diploidization, is characteristic of the mesohexaploid taxa (46 - 73% genome diploidization) and the mesotetraploid Australian Microlepidieae (66%). This pattern may be explained by the age of mesohexaploidies, whereby a two-step hybridization event resulting in a hexaploid genome theoretically needs more time than a one-step tetraploidization, and/or by higher levels of genome duplication (6*x* vs. 4*x*) increasing the frequency of inter-genome recombination and, at the same time, buffering potentially deleterious consequences of chromosomal rearrangements. The least diploidized genomes were found among Thelypodieae and *Pachycladon* species (14 and 37%, respectively), i.e. two mesopolyploid groups which originated either relatively recently (Thelypodieae) or with a slow genome diploidization (*Pachycladon*).

**Figure 5.**
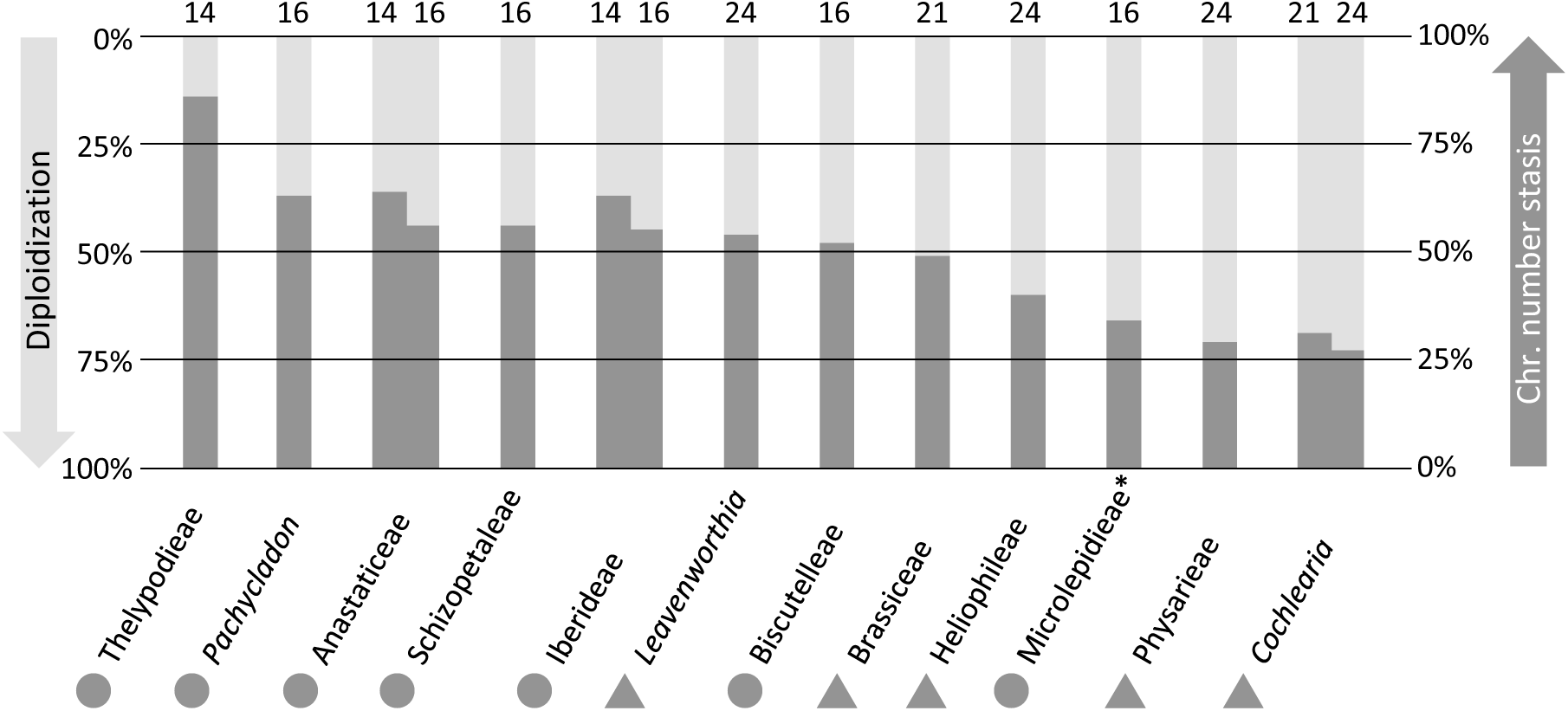
Relative percentage of post-polyploid diploidization and ancestral chromosome number stasis in 12 mesopolyploid crucifer taxa. The percentage of diploidization and chromosome number stasis, respectively, corresponds to the ratio between an average extant chromosome number and the inferred ancestral chromosome number (e.g., if the average extant chromosome number equal 6.5 and the ancestral number is 24, the level of diploidization equal 73%). Numbers above the graph denote the inferred ancestral mesopolyploid chromosome number for a given taxon; circular and triangular symbols refer to mesotetraploid and mesohexaploid events, respectively. Two possible ancestral chromosome numbers are given for Anastaticeae, Cochlearieae and Iberideae. As the diploidization tempo is strikingly different for *Pachycladon* and the remaining Microlepidieae genera (*), these groups are shown separately.

### Gene retention and loss following multiple, independent mesopolyploidies

Despite substantial cytological variation across the nine independent mesopolyploidies studied in our analyses, we observed a convergent pattern among the GO categories of genes retained and lost. Eight of the nine independent mesopolyploidies had similar patterns of GO category retention among their paralogs following WGD. The pattern of the other mesopolyploidy was similar to the GO patterns of genes retained in duplicate following the At-*α* event. The overall pattern of convergent, biased GO category enrichment following WGD across the nine mesopolyploidies was consistent with the predictions of the dosage balance hypothesis (DBH). The DBH predicts that genes with many connections or in dosage sensitive pathways will be more highly retained following polyploidy (Freeling 2009; Birchler and Veitia 2007, 2010, 2012; Edger and Pires 2009). The higher than expected retention is proposed to maintain the stoichiometry and “balance” of these biological pathways. Similarly, the same genes will be lost at higher rate from small scale duplications to minimize dosage problems. Among the many hypotheses for duplicate gene retention (Kondrashov and Kondrashov 2006; Freeling 2009), the DBH is the only hypothesis that explicitly predicts the parallel retention and loss of functionally related genes across species following WGD (Freeling and Thomas 2006; Freeling 2009; Conant *et al*. 2014). Previous research has found evidence consistent with the DBH from analyses of gene ontology (GO) enrichment among the genes duplicated by paleopolyploidy, including the At-*α* WGD in the Brassicaceae (Maere *et al*. 2005; Thomas *et al*. 2006; Bekaert *et al*. 2011; Conant *et al*. 2014) and other plant WGDs (Rensing *et al*. 2007; Barker *et al*. 2008; Shi *et al*. 2010; Geiser *et al*. 2016).

Consistent with the DBH, we also found parallel patterns of gene retention and loss among the GO categories of At-*α* paralogs. Notably, the patterns of GO category retention across the mesopolyploid and At-*α* paralogs exhibit some differences. Previous analyses have found differences in the patterns of gene retention and loss across WGDs of different ages in the Brassicaceae (Maere *et al*. 2005; Bekaert *et al*. 2011) and the Actinidiaceae (Shi *et al*. 2010). Although we may expect little difference in the pattern of gene retention and loss from any WGD under the DBH, as observed in some analyses (e.g., Barker et al. 2008), dosage sensitivity appears to be fluid and may change over time (Conant *et al*. 2014). Thus, we may observe variation in the functional patterns of gene retention and loss across WGDs of different ages.

Our results extend past analyses by demonstrating convergent biased gene retention and loss following numerous independent polyploidies. Previous analyses of gene retention and loss for multiple species found parallel patterns of gene retention and loss across multiple WGDs in the Asteraceae (Barker *et al*. 2008), including rapid emergence of this pattern in only 40 generation old *Tragopogon* polyploids (Buggs *et al*. 2012). Thus, our observation of convergent gene retention and loss across multiple mesopolyploidies of the Brassicaceae is a new but expected result. The DBH is the most likely explanation for the parallel retention as it is difficult to conceive that sub- or neofunctionalization would consistently yield the same pattern of paralog retention across nine independent polyploidies in 13 different species. Further, our chromosome painting analyses uncovered significant differences in the degree of chromosomal and genome-wide reorganization and diploidization. Despite these differences, we still found that these genomes retain similar GO categories of genes across their independent WGDs as expected by the DBH.

### Mesopolyploidy and diversification

Polyploidization is frequently linked with adaptive radiation and diversification (Schranz *et al*. 2012; Vanneste *et al*. 2014; Tank *et al*. 2015; Shimizu-Inatsugi *et al*. 2016; Soltis and Soltis 2016). However, establishing a causal link between mesopolyploidies and diversification is not straightforward, as several intrinsic and extrinsic factors have to be considered (e.g., age of a WGD, geographical and paleoclimatic constraints, frequency of neopolyploid formation, extinction rate). In Brassicaceae, independent WGDs appear to pre-date the origins of some species-rich groups, such as Brassiceae (47 genera: 229 species), Heliophileae (2: 100), Physarieae (7: 135) and Thelypodieae (28: 249), and moderately large groups such as Anastaticeae (13: 74) and the Microlepidieae (16: 56). This pattern is however not universal as mesopolyploidy also occurs in the ancestry of relatively small genera and tribes, such as Biscutelleae (2 genera: 45 species), Cochlearieae (2: 29), Iberideae (2: 30), Schizopetaleae (3: 16) and *Leavenworthia* (8 spp.). Among 11 Brassicaceae tribes harboring more than 100 species each (Al-Shehbaz 2012), most likely only three (Brassiceae, Physarieae and Thelypodieae) have a mesopolyploid origin, whereas the species richness of the remaining seven tribes (Alysseae, Arabideae, Boechereae, Cardamineae, Erysimeae, Euclidieae, and Lepidieae) is explained by neopolyploidy (Hohmann *et al*. 2015). In the light of our results, we analyzed the percentage of neopolyploids within mesopolyploid clades and concluded that with the exception of *Cochlearia* (82%), neopolyploidy maximally explains 25% of species richness of these clades **(Fig. S2)**. Thus, a preliminary conclusion can be drawn that Brassicaceae mesopolyploids diversify by independent diploidizations presumably associated with reproductive isolation and speciation instead of subsequent rounds of (neo) polyploidization.

## Experimental procedures

### Plant material

Seeds of each species were donated by botanical gardens and individuals, or collected in the field **(Table S1)**. All plants were grown from seed in a greenhouse or growth chamber until flowering.

### Chromosome preparation and chromosome counts

In preparation for comparative chromosome painting (CCP), entire inflorescences were fixed in the ethanol: acetic acid fixative (3:1) overnight and stored in 70% ethanol at −20°C. Selected inflorescences or individual flower buds were rinsed and digested as described by Lysak and Mandáková (2013); the same protocol was applied to prepare chromosome spreads from individual flower buds. Suitable slides staged with a phase contrast microscope were used for CCP experiments as well as to establish chromosome numbers from mitoses of tapetal tissues. Prior to CCP, slides were pretreated by pepsin and RNase (Lysak and Mandáková 2013). A high amount of cytoplasm covering pachytene chromosomes in *Heliophila africana, Streptanthus farnthworthianus* and *Schizopetalon walkeri* required twice as long of a pepsin treatment compared to the other species. Chromosome numbers were recorded from microscope photographs after counterstaining with DAPI (4’,6-diamidino-2-phenylindole; 2 mg/mL) in Vectashield.

### Comparative chromosome painting

Preparation of painting probes and their localization on chromosomes followed established protocols (Lysak and Mandáková 2013). Briefly, chromosome-specific BAC contigs of *A. thaliana*, corresponding to 12 GBs that comprise four ancestral crucifer chromosomes (see Fig. 1 and Table 2 in Lysak *et al*. 2016), were used as painting probes. DNA of individual BAC clones was isolated by a standard alkaline lysis protocol and labeled with biotin-, digoxigenin-, and Cy3-dUTP by nick translation. Individually labeled probes were pooled according to the 12 GBs, ethanol precipitated, dissolved in hybridization buffer, and hybridized to pepsin- and RNase treated chromosome preparations for c. 60 hrs (for details see Lysak and Mandáková 2013). The concentration of painting probes was 200 ng/BAC/slide for mesotetraploid species (except for a double concentration of 400 ng/BAC/slide in *Lobularia libyca*) and 300 ng/BAC/slide for mesohexaploid species. After the hybridization biotin-and digoxigenin-labeled painting probes were immunodetected using fluorescently labeled antibodies, dehydrated and DAPI-stained. Painted chromosomes were photographed using an Olympus BX-61 epifluorescence microscope equipped with a Zeiss CoolCube camera. Monochromatic images were pseudocolored and merged using the Adobe Photoshop CS5 software.

### Transcriptome assembly

RNA was isolated from fresh young leaves using the RNeasy plant mini kit (Qiagen). Raw reads for each species were demultiplexed, cleaned with SnoWhite (Dlugosch *et al*. 2013), and assembled into contigs **(Table S2)**. Two different assembly strategies were used for our two different types of sequence data. SOAPdenovo-Trans (Xie *et al*. 2014) was used to assemble the paired end Illumina reads using a k-mer of k = 57. All other parameters were set to default. For our single Ion Torrent transcriptome, we treated the data similarly to 454 reads and assembled them with MIRA version 3.2.1 (Chevreux *et al*. 2004) using the ‘accurate.est.denovo.454’ assembly mode. As MIRA may split up high coverage contigs into multiple contigs, we used CAP3 at 94% identity to further assemble the MIRA contigs and singletons (Huang and Madan 1999).

### DupPipe analyses of WGDs from transcriptomes of single species

For each transcriptome, we used our DupPipe pipeline to construct gene families and estimate the age of gene duplications (Barker *et al*. 2008, 2010). We translated DNA sequences and identified reading frames by comparing the Genewise alignment to the best hit protein from a collection of proteins from 25 plant genomes from Phytozome (Goodstein *et al*. 2012). For all DupPipe runs, we used protein-guided DNA alignments to align our nucleic acids while maintaining reading frame. We estimated synonymous divergence (Ks) using PAML with the F3×4 model (Yang 2007) for each node in our gene family phylogenies. We identified peaks of gene duplication as evidence of ancient WGDs in histograms of the age distribution of gene duplications (Ks plots). We used a mixture model, EMMIX (McLachlan *et al*. 1999), to identify significant peaks and estimate their median Ks values. Significant peaks were identified by comparing the Δ BIC values for models with and without inferred polyploid peaks. We used the Bayesian Information Criterion (BIC) instead of the Akaike Information Criterion (AIC) because the BIC has more severe penalties for increasing parameters (peaks, in this application).

### Estimating orthologous divergence

To estimate the average ortholog divergence of conifer taxa and compare to observed paleopolyploid peaks, we used our previously described RBH Ortholog pipeline (Barker *et al*. 2010). Briefly, we identified orthologs as reciprocal best blast hits in pairs of transcriptomes. Using protein-guided DNA alignments, we estimated the pairwise synonymous (Ks) divergence for each pair of orthologs using PAML with the F3×4 model (Yang 2007). We plotted the distribution of ortholog divergences and calculated the median divergence to compare against the synonymous divergence of paralogs from inferred WGDs in related lineages. For *Pachycladon* and *Stenopetalum* we also used the orthologs to assess substitution rate heterogeneity (see Methods S1).

### MAPS analyses of WGDs from transcriptomes of multiple species

To further infer and locate paleopolyploidy in our data sets, we used our recently developed gene tree sorting and counting algorithm, the Multi-tAxon Paleopolyploidy Search (MAPS) tool (Li *et al*. 2015). The species trees for MAPS analyses were based on previously published phylogenies (Huang *et al*. 2016). We circumscribed and constructed nuclear gene family phylogenies from multiple species for the MAPS analyses. For further information on the MAPS analyses see Methods S2.

### Gene Ontology (GO) annotations and paleolog retention and lost patterns

Gene Ontology (GO) annotations of Brassicaceae all transcriptomes and paleologs were obtained through discontiguous MegaBlast searches against annotated *Arabidopsis thaliana* transcripts from TAIR (Swarbreck *et al*. 2008) to find the best hit with length of at least 100 bp and an e-value of at least 0.01. For each species, we assigned paralogs to the independent mesopolyploidies and *At-α* WGD based on the Ks ranges identified in mixture model analyses (McLachlan *et al*. 1999; Barker *et al*. 2008). Further information on the GO analysis can be found in Methods S3.

## Supporting Information

Additional supporting information may be found in the online version of this article.

**Figure S1** Phylogenetic scheme of the Brassicaceae with a tentative placement of 13 known mesotetraploid (white stars) and mesohexaploid (black stars) events discussed in the paper.

**Figure S2** Relationship between species richness and percentage of neopolyploid species for 11 mesopolyploid crucifer taxa.

**Table S1** Origin and collection data for species used in the present study.

**Table S2** Assembly statistics for 13 transcriptomes used in this study.

**Table S3** Sampling and summary statistics for MAPS analyses.

**Data S1** Patterns of chromosomal homoeology revealed by comparative chromosome painting (CCP) in ten mesopolyploid Brassicaceae species.

**Data S2** Mesopolyploid Brassicaceae clades, diversification and colonization.

**Methods S1** Estimating substitution rate heterogeneity.

**Methods S2** Multi-tAxon Paleopolyploidy Search (MAPS) analysis.

**Methods S3** Gene Ontology annotations.

## Acknowledgments

The work was supported by the Czech Science Foundation (grant 13-10159S) and the CEITEC 2020 (LQ1601) project to T.M. and M.L., and by NSF-IOS-1339156 and NSF-EF-1550838 to M.S.B. and Z.L. We thank J. Busch, A. Doust, P. B. Heenan, Ch. Konig, and Millenium Seed Bank, Royal Botanic Gardens, Kew for providing seeds of species used in this study.

